# *Drosophila* phosphatidylinositol-4 kinase *fwd* promotes mitochondrial fission and can suppress *Pink1/parkin* phenotypes

**DOI:** 10.1101/2020.05.13.093823

**Authors:** Ana Terriente-Felix, Emma L. Wilson, Alexander J. Whitworth

## Abstract

Balanced mitochondrial fission and fusion play an important role in shaping and distributing mitochondria, as well as contributing to mitochondrial homeostasis and adaptation to stress. In particular, mitochondrial fission is required to facilitate degradation of damaged or dysfunctional units via mitophagy. Two Parkinson’s disease factors, PINK1 and Parkin, are considered key mediators of damage-induced mitophagy, and promoting mitochondrial fission is sufficient to suppress the pathological phenotypes in *Pink1/parkin* mutant *Drosophila*. We sought additional factors that impinge on mitochondrial dynamics and which may also suppress *Pink1/parkin* phenotypes. We found that the *Drosophila* phosphatidylinositol 4-kinase IIIβ homologue, Four wheel drive (Fwd), promotes mitochondrial fission downstream of the pro-fission factor Drp1. Previously described only as male sterile, we identified several new phenotypes in *fwd* mutants, including locomotor deficits and shortened lifespan, which are accompanied by mitochondrial dysfunction. Finally, we found that *fwd* overexpression can suppress locomotor deficits and mitochondrial disruption in *Pink1/parkin* mutants, consistent with its function in promoting mitochondrial fission. Together these results shed light on the complex mechanisms of mitochondrial fission and further underscore the potential of modulating mitochondrial fission/fusion dynamics in the context of neurodegeneration.

**Author Summary:** Mitochondria are dynamic oganelles that can fuse and divide, in part to facilitate turnover of damaged components. These processes are essential to maintain a healthy mitochondrial network, and, in turn, maintain cell viability. This is critically important in high-energy, post-mitotic tissues such as neurons. We previously identified *Drosophila* phosphatidylinositol-4 kinase *fwd* as a pro-fission factor in a cell-based screen. Here we show that loss of *fwd* regulates mitochondrial fission *in vivo*, and acts genetically downstream of *Drp1*. We identified new phenotypes in *fwd* mutants, similar to loss of *Pink1/parkin*, two genes linked to Parkinson’s disease and key regulators of mitochondrial homeostasis. Importantly, *fwd* overexpression is able to substantially suppress locomotor and mitochondrial phenotypes in *Pink1/parkin* mutants, suggesting manipulating phophoinositides may represent a novel route to tackling Parkinson’s disease.

## Introduction

Mitochondria are dynamic organelles that are transported to the extremities of the cell and frequently undergo fusion and fission events, which influences their size, branching and degradation. Many of the core components of the mitochondrial fission and fusion machineries have been well characterised, these include the pro-fusion factors Mfn1/2 and Opa1, and pro-fission factors Drp1 and Mff (1). Maintaining an appropriate balance of fission and fusion, as well as transport dynamics, is crucial for cellular health and survival as mutations in many of the core components cause severe neurological conditions in humans and model organisms (2).

The mitochondrial fission/fusion cycle has been linked to the selective removal of damaged mitochondria through the process of autophagy (termed mitophagy), in which defective mitochondria are engulfed into autophagosomes and degraded by lysosomes (3, 4). Two genes that have been firmly linked to the mitophagy process are *PINK1* and *PRKN* (5-7). Mutations in these genes cause autosomal-recessive juvenile parkinsonism, associated with degeneration of midbrain dopaminergic neurons and motor impairments, among other symptoms and pathologies. Studies from a wide variety of model systems have shown various degrees of mitochondrial dysfunction associated with mutation of *PINK1/PRKN* homologues including disrupted fission/fusion (8-16). *Drosophila* have proven to be a fruitful model for investigating the function of the conserved homologues *Pink1* and *parkin*, with these mutants exhibiting robust mitochondrial disruption and neuromuscular phenotypes. Importantly, several studies have shown that the pathological consequences of loss of *Pink1* or *parkin* can be largely suppressed by genetic manipulations that increase mitochondrial fission or reduce fusion (17-23).

To identify genes involved in mitochondrial quality control and homeostasis, we previously performed an RNAi screen in *Drosophila* S2 cells to identify kinases and phosphatases that phenocopy or suppress hyperfused mitochondria caused by loss of *Pink1* (24). We identified the phosphatidylinositol 4-kinase IIIβ homologue, *four wheel drive* (*fwd*), whose knockdown phenocopied *Pink1* RNAi, resulting in excess mitochondrial fusion. *Drosophila* mutant for *fwd* have been reported to be viable but male sterile due to incomplete cytokinesis during spermatogenesis (25-28); however, no other organismal phenotypes or mitochondrial involvement have been described to date. Thus, we sought to better understand the role of Fwd in mitochondrial homeostasis.

In this study, we have characterised *fwd* mutants for organismal phenotypes associated with *Pink1/parkin* dysfunction, and analysed the impact on mitochondrial form and function. We have also investigated genetic interactions between *fwd* and *Pink1/parkin*, as well as with mitochondrial fission/fusion factors. We found that loss of *fwd* inhibited mitochondrial function, causing increased mitochondrial length and branching, and decreased respiratory capacity. These effects were associated with shortened lifespan and dramatically reduced locomotor ability, similar to *Pink1* and *parkin* mutants. Furthermore, *fwd* overexpression was sufficient to significantly suppress *Pink1/parkin* mutant locomotor deficits and mitochondrial phenotypes. Interestingly, we found that the mitochondrial and locomotion phenotypes in *fwd* mutants can be rescued by loss of pro-fusion factors *Marf* and *Opa1*, but the activity of *Drp1* appears to require *fwd*. These results support a role for *fwd* in regulating mitochondrial morphology, specifically in facilitating mitochondrial fission, and further substantiate the important contribution of aberrant mitochondrial fission/fusion dynamics in *Pink1/parkin* phenotypes.

## Results

### Loss of *fwd* causes mitochondrial hyperfusion along with locomotor and lifespan deficits

We previously found that knockdown of *fwd* phenocopied loss of *Pink1* in cultured cells by causing mitochondrial hyperfusion (24). To extend these *in vitro* observations we sought to determine whether *fwd* has a broader role in regulating mitochondrial homeostasis *in vivo*. In striking similarity to *Pink1* mutants, mutations in *fwd* have previously been shown to cause male sterility due to aberrant spermatogenesis (25-28); however, no other organismal phenotypes have been described.

*Pink1* mutants have a range of additional phenotypes including deficits in negative geotaxis (climbing ability), disruption of flight muscle mitochondria, shortened lifespan, and modest degeneration of dopaminergic (DA) neurons (10, 29). Thus, we assessed these phenotypes in two *fwd* mutants – a nonsense mutation, *fwd*^3^, and a *P*-element insertion, *fwd*^neo1^. In all instances, these mutations were crossed to a deficiency (*Df(3L)7C*) to avoid potential extragenic effects from homozygosity. Both mutant combinations, *fwd*^3^/Df and *fwd*^neo1^/Df (hereafter, designated simply as *fwd*^3^ and *fwd*^neo1^), displayed a striking loss of climbing ability in young flies (Fig. 1A), though the phenotype was weaker in *fwd*^neo1^ consistent with it being a hypomorph. Notably, transgenic re-expression of *fwd* using a ubiquitous driver (*da*-*GAL4*), was able to restore climbing ability to near wild-type levels (Fig. 1A), supporting the specificity of this phenotype for loss of *fwd*. Analysing longevity in the *fwd*^3^ null mutants, revealed a significant reduction in median lifespan (Fig. 1B). However, no significant loss of DA neurons was detected in aged *fwd* mutant brains (Fig. 1C). These results reveal some phenotypic similarity between *Pink1* and *fwd* mutants at the organismal level as well as the cellular level.

**Figure 1.**
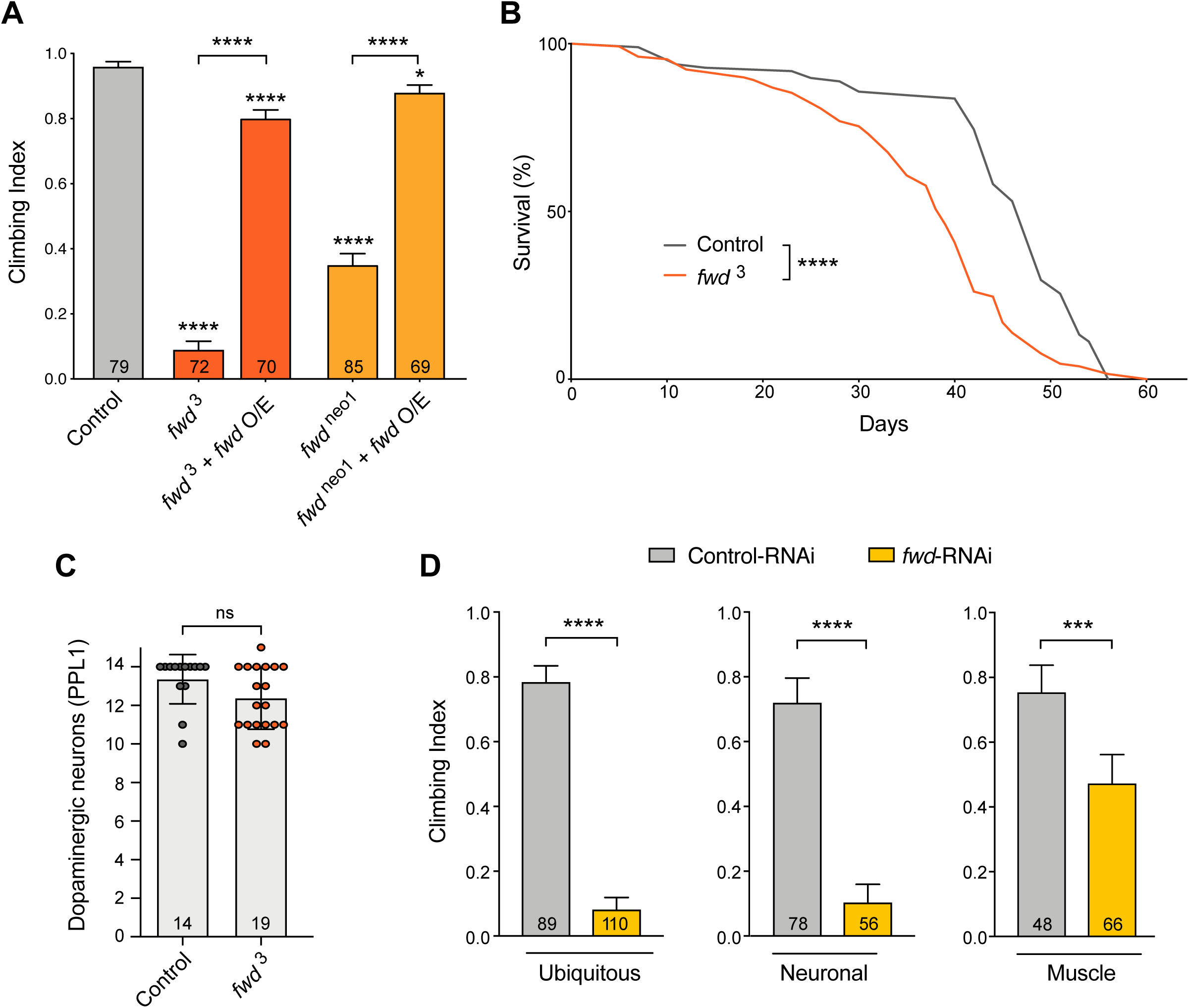
Loss of *fwd* causes motor deficits and shortened lifespan. (A) Climbing assay of *fwd* mutants (*fwd*^3^ and *fwd*^neo1^) in trans to a deficiency (Df), alone or with transgenic overexpression of *fwd* (*fwd* O/E) driven by *da*-*GAL4*. (B) Lifespan analysis of control and *fwd* mutants. Significance for lifespan was analysed by log-rank (Mantel-Cox) test. (C) Quantification of dopaminergic neurons in PPL1 cluster of 30-day-old adult brains. Chart shows mean ± SD with individual data points. Significance was analysed by Mann-Whitney test. (D) Climbing analysis of *fwd* knockdown (RNAi) in all (ubiquitous) or selected tissues. Charts show mean ± 95% confidence interval (CI); number of animals analysed is shown in each bar. Significance for climbing was analysed by Kruskal-Wallis test with Dunn’s post-hoc correction for multiple comparisons. Comparison is against the control unless otherwise indicated; * *P<*0.05, *** *P<*0.001, **** *P<*0.0001; ns, non-significant. Full genotypes are given in Supplementary Table 1.

To investigate the relative contribution of *fwd* to locomotor ability in different tissues, we expressed a transgenic RNAi construct (30) via tissue-specific drivers. We first verified that ubiquitous knockdown of *fwd* via *da*-*GAL4* phenocopied the genetic mutants, thus, demonstrating its efficacy to recapitulate null mutant phenotypes (Fig. 1D). Interestingly, pan-neuronal knockdown, using *nSyb-GAL4*, reproduced the striking loss of climbing ability, whereas knockdown in all muscles via *Mef2-GAL4* only modestly affected climbing (Fig. 1D). Thus, *fwd* shows some tissue-selective requirement but plays an important role in the nervous system that was not previously appreciated. Since *fwd* knockdown in cultured cells caused mitochondrial fusion, similar to loss of *Pink1*, we sought to further characterise the impact of *fwd* loss on mitochondria *in vivo*. Mitochondria are particularly abundant in adult flight muscles, and this tissue is severely affected in *Pink1/parkin* mutants (8, 10, 29), so we first analysed mitochondrial morphology in *fwd* mutants in this tissue. Imaging mitochondria by fluorescence or electron-microscopy in flight muscles revealed them to be grossly normal in their cristae structure, size and abundance compared to control (Fig. 2A-B).

**Figure 2.**
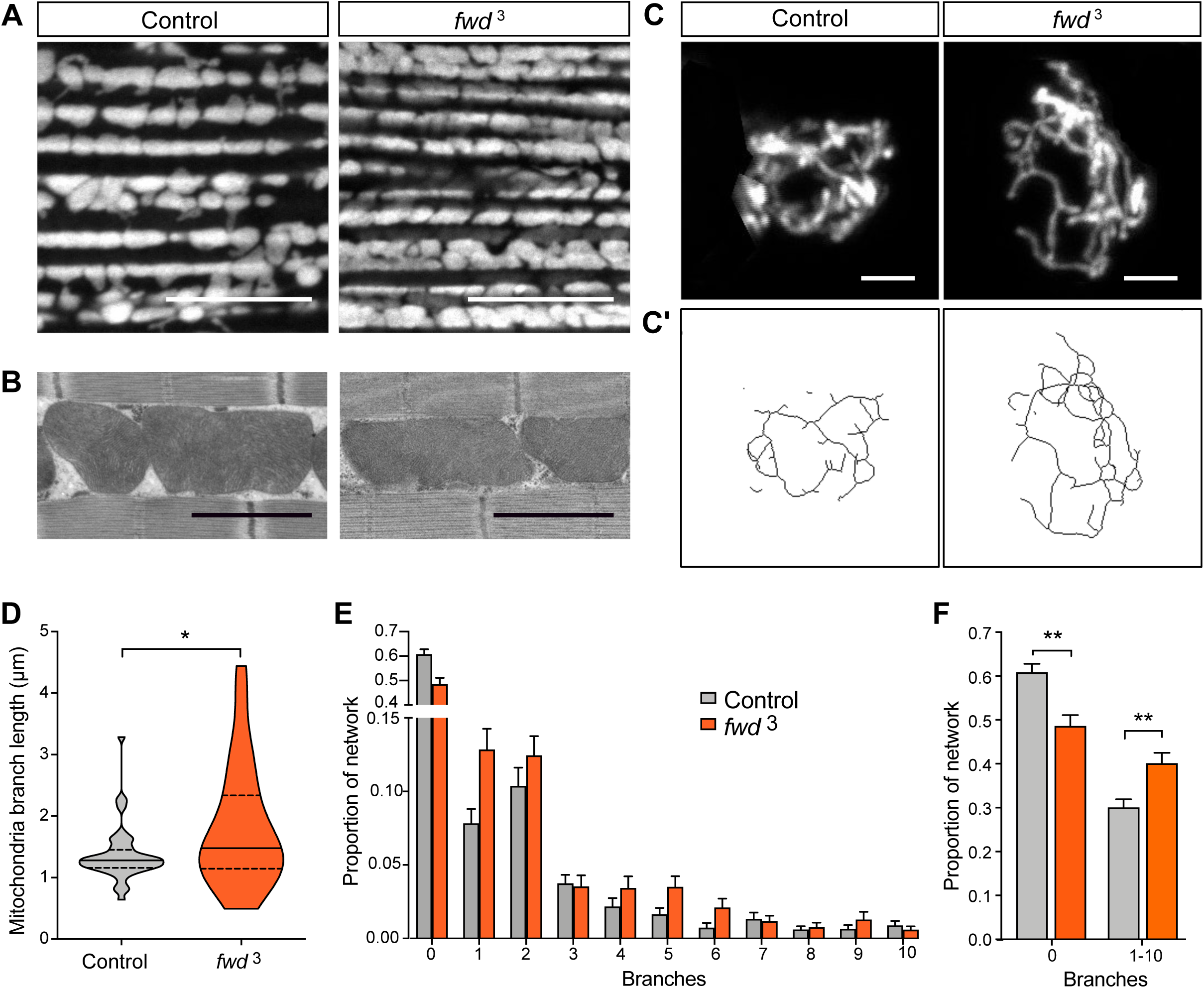
Loss of *fwd* causes excess mitochondrial fusion. (A) Confocal microscopy analysis of mitochondrial morphology, visualised using mitoGFP, in control and *fwd* mutant flight muscles. Scale bar = 10 μm. (B) Electron-microscopy analysis of mitochondrial structure in flight muscles. Scale bar = 1 μm. (C) Confocal microscopy analysis of mitochondrial network morphology (mitoGFP) in neuronal cell bodies from larval ventral ganglion of control and *fwd* mutants. Image shows a projected z-stack. Scale bar = 2 μm. (C’) Skeletonised image of mitochondrial network used for quantification. (D) Quantification of median mitochondrial branch length per cell. Violin plot indicating median (thick, horizontal line) and quartiles (dashed lines). Significance was analysed by Mann-Whitney test. (E) Frequency distribution plot of mitochondrial network connectivity (number of branches) per cell. N = 46 (control) and 54 (*fwd*). (F) Chart summarising quantification of connectivity shown in E, plotting the proportion of individual mitochondria (0 branches) and connected mitochondria (1-10 branches) relative to the total number of networks per cell. Significance was analysed by Kruskal-Wallis test with Dunn’s post-hoc correction for multiple comparisons. * *P<*0.05, ** *P<*0.01. Full genotypes are given in Supplementary Table 1.

We next sought to analyse the mitochondrial morphology in a tissue where the specific knockdown of *fwd* resulted in strong climbing defects. We analysed the network morphology in cell bodies of the larval ventral ganglion (part of the central nervous system). Expression of mitoGFP in a subset of neurons, driven by *CCAP*-*GAL4*, allowed better three-dimensional imaging of the mitochondrial network (Fig. 2C). While the overall appearance was similar between *fwd* mutant and control, quantitative analysis of the networks revealed that both the length and connectivity (number of branches) were increased upon loss of *fwd* (Fig. 2D-F). These results are consistent with the previous cell-based study indicating loss of *fwd* causes mitochondrial hyperfusion.

We next assessed mitochondrial function, analysing maximal respiratory capacity in intact mitochondria and overall ATP levels in whole animals. Respiration measured by the oxygen consumption rate in energised mitochondria, stimulated via either complex I or complex II substrates, was significantly reduced in *fwd* mutants (Fig. 3A). However, the overall level of ATP was not significantly affected (Fig. 3B). These results indicate that mitochondrial respiration is affected by loss of *fwd* but compensatory mechanisms could still maintain normal steady-state ATP levels in the organism.

**Figure 3.**
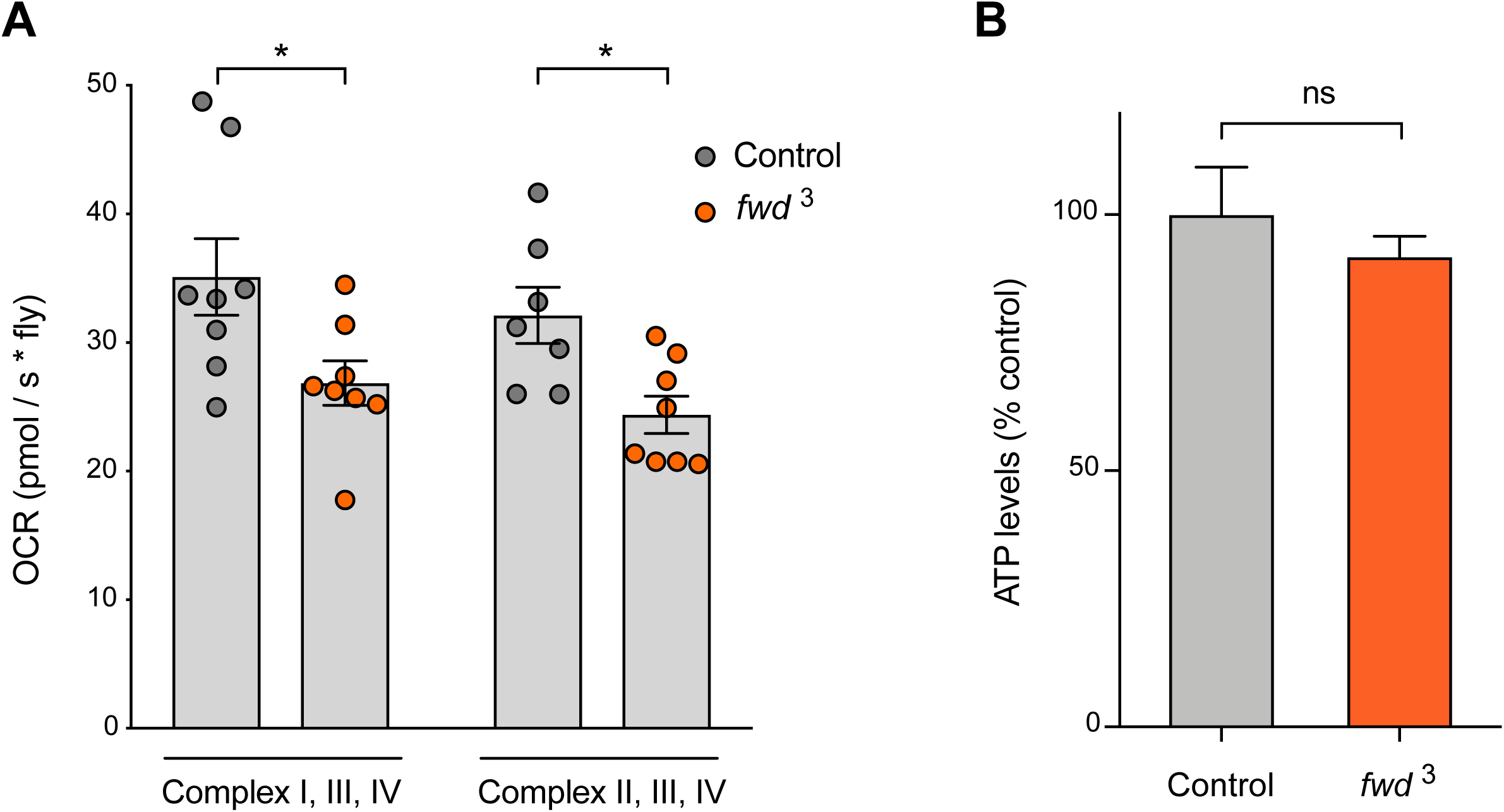
Loss of *fwd* inhibits mitochondrial respiration. (A) Mitochondrial respiration analysis by oxygen consumption rate (OCR) and (B) ATP levels in control and *fwd* mutant adults. Charts show mean ± SEM. Significance was analysed by paired (A) or unpaired (B) *t*-test. * *P<*0.05; ns, non-significant. Full genotypes are given in Supplementary Table 1.

### *fwd* mutant phenotypes are suppressed by loss of fusion factors

The results above substantiate that loss of *fwd* causes excess mitochondrial fusion *in vivo*. We next addressed whether the mitochondrial hyperfusion may contribute to the locomotor deficit. To do this we combined ubiquitous expression of *fwd* RNAi with genetic manipulations that reduce fusion (partial loss of pro-fusion factors *Marf* or *Opa1*) or promote fission (overexpression of pro-fission factor *Drp1*), and assessed climbing behaviour. Heterozygous loss of either *Marf* (the fly homologue of *MFN1/2*) or *Opa1*, which did not affect climbing alone, was sufficient to significantly suppress the climbing deficit caused by *fwd* RNAi (Fig. 4A, B). However, contrary to what we expected, overexpression of *Drp1* was not able to ameliorate the climbing defect (Fig. 4C).

**Figure 4.**
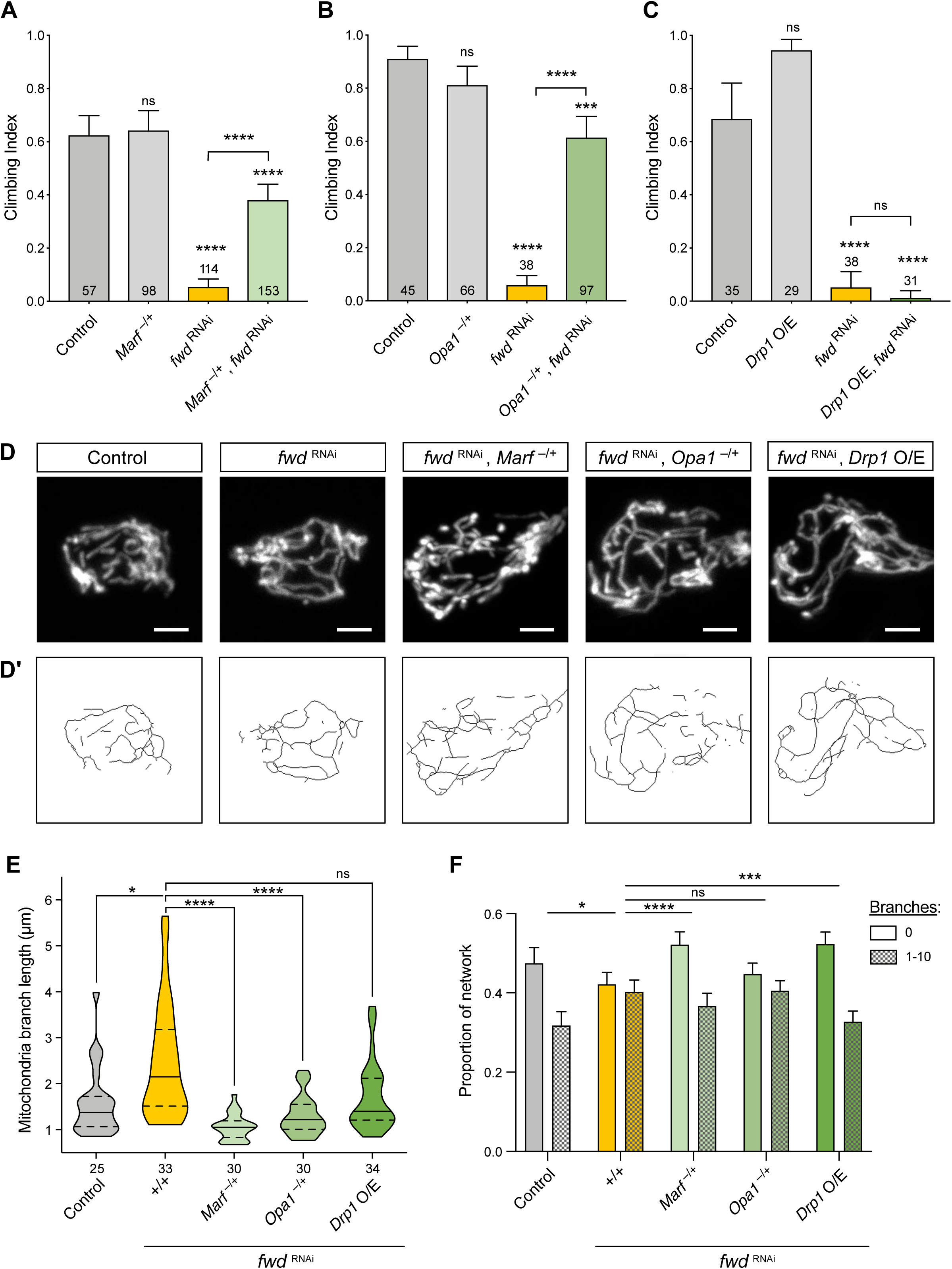
*fwd* genetically interacts with mitochondrial fission/fusion factors. (A-C) Climbing assay of *fwd* RNAi alone or in combination with heterozygous *Marf* or *Opa1* mutations or transgenic overexpression of *Drp1*. Transgenic expression was mediated via *da-GAL4*. Charts show mean ± 95% confidence interval (CI); number of animals analysed is shown in each bar. (D) Confocal microscopy analysis of mitochondrial network morphology (mitoGFP) in neuronal cell bodies from larval ventral ganglion of control, *fwd* RNAi alone and in combination with heterozygous *Marf* or *Opa1* mutations or transgenic overexpression of *Drp1*. Image shows a projected z-stack. Scale bar = 2 μm. (D’) Skeletonised image of mitochondrial network used for quantification. (E) Quantification of median mitochondrial branch length per cell. Violin plot indicating median (thick, horizontal line) and quartiles (dashed lines). Significance was analysed by Mann-Whitney test. (F) Plot of the proportion of individual mitochondria (0 branches) and connected mitochondria (1-10 branches) quantified per cell. Significance was calculated by Kruskal-Wallis test with Dunn’s post- hoc correction for multiple comparisons. Comparison is against the control unless otherwise indicated; * *P<*0.05, *** *P<*0.001, **** *P<*0.0001; ns, non-significant. Full genotypes are given in Supplementary Table 1.

To better understand these results, we analysed the mitochondrial morphology in neuronal cell bodies of these genotypes. As with the *fwd* mutant, *fwd* RNAi caused a significant elongation of mitochondria and increased branching (Fig. 4D-F). Consistent with the effects on climbing, heterozygous loss of *Marf* or *Opa1* reverted the increase in mitochondrial length, whereas *Drp1* overexpression did not (Fig. 4D, E). Interestingly, the increased branching caused by loss of *fwd* was suppressed by heterozygous loss of *Marf* or *Drp1* overexpression, but not by heterozygous loss of *Opa1* (Fig. 4D, F). The reasons for the complex effects on branching are unclear but may reflect that Marf directs fusion of the OMM (and hence, coordinates branching), while Opa1 regulates fusion of IMM. Nevertheless, the effects on mitochondrial branch length suggest that Drp1 may require Fwd to execute mitochondrial fission. Overall, the genetic interaction of *Marf* and *Opa1* suppressing the *fwd* RNAi-induced climbing deficit supports this phenotype being, at least partially, caused by mitochondrial hyperfusion.

### *fwd* overexpression can suppress *Pink1/parkin* mutant phenotypes

While many studies have focused on the role of PINK1/Parkin in damage-induced mitophagy, aberrant mitochondrial dynamics is clearly a major cause of *Pink1/parkin* mutant phenotypes in *Drosophila*, including locomotor deficits and flight muscle degeneration, since these can be substantially suppressed by promoting mitochondrial fission (17-20). As our results indicate that Fwd promotes mitochondrial fission, we next tested whether overexpression of *fwd* could ameliorate *Pink1* and *parkin* mutant phenotypes. Combining *Pink1/parkin* mutants with ubiquitous *fwd* overexpression was sufficient to significantly suppress the climbing deficit in both mutants (Fig. 5A, B). In addition, the thoracic indentations caused by degeneration of the underlying flight muscle were also significantly improved (Fig. 5C). Disruption of mitochondrial integrity in the flight muscles was also visibly improved when *fwd* was overexpressed in muscles (Fig. 5D). These results are consistent with Fwd overexpression promoting mitochondrial fission and partially reverting the hyperfusion caused by *Pink1/parkin* loss.

**Figure 5.**
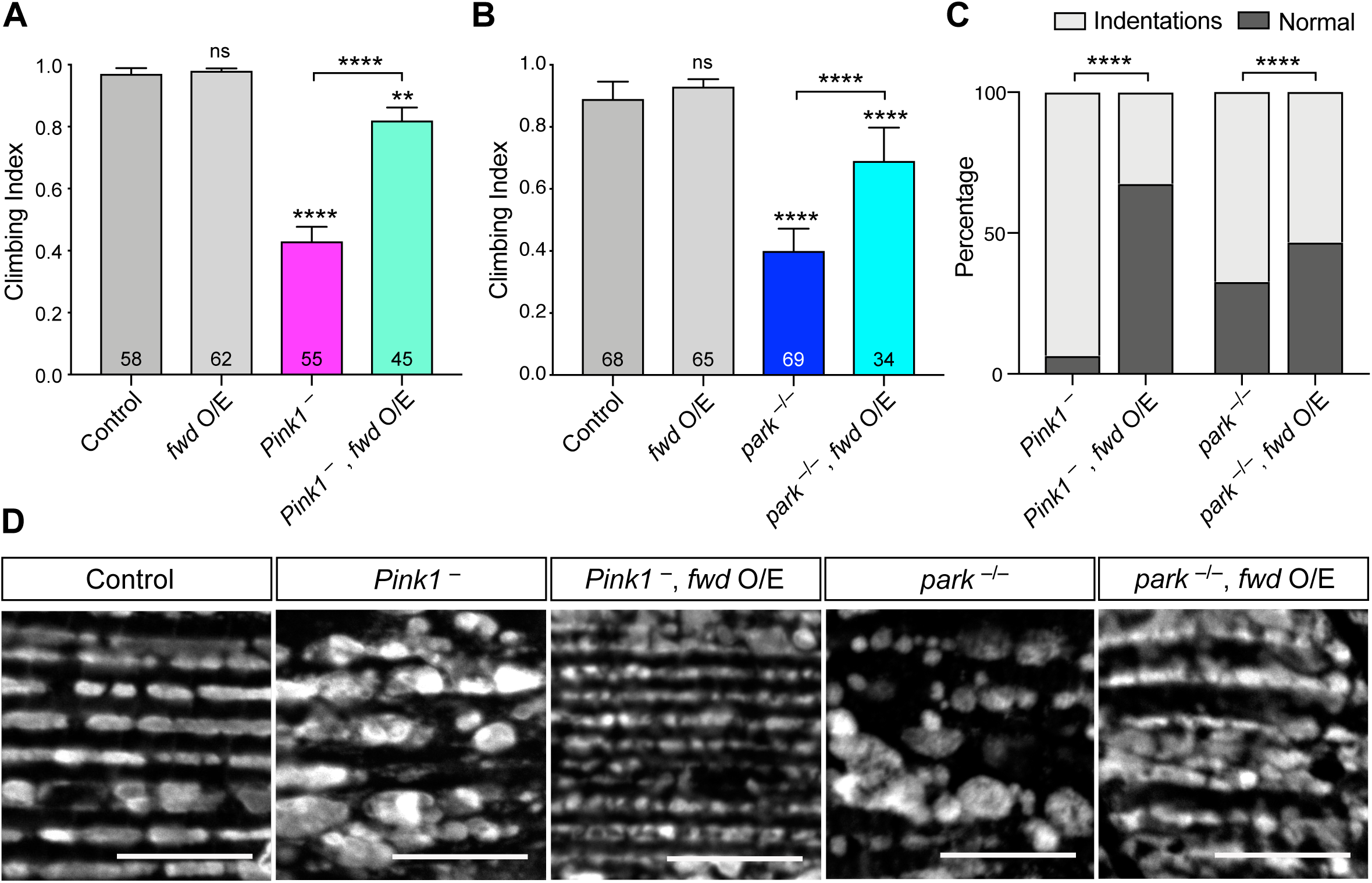
*fwd* overexpression partially suppresses *Pink1/parkin* phenotypes. (A, B) Climbing assay of control, *Pink1* or *parkin* mutants with or without *fwd* overexpression induced by *da-GAL4*. Significance was analysed by Kruskal-Wallis test with Dunn’s post-hoc correction for multiple comparisons. Comparison is against the control unless otherwise indicated; * *P<*0.05, *** *P<*0.001, **** *P<*0.0001; ns, non-significant. (C) Analysis of thoracic indentations evident in *Pink1* or *parkin* mutants in the presence of absence of *fwd* overexpression, induced by *da-GAL4*. Significance was determined by Chi-squared test. **** *P<*0.0001. (D) Confocal microscopy analysis of mitochondrial integrity, visualised by anti-ATP5A immunostaining, in flight muscles of the indicated genotypes. Transgenic expression was mediated via *Mef2-GAL4*. Scale bar = 10 μm. Full genotypes are given in Supplementary Table 1.

We were intrigued by the earlier observation that heterozygous loss of *Marf* or *Opa1* could revert the aberrant mitochondrial morphology and climbing defect of *fwd* RNAi, but the overexpression of *Drp1* did not (Fig. 4). These results suggested that the activity of Drp1 might require Fwd, which we sought to test further. As a paradigm for Drp1 activity, overexpression of *Drp1* is sufficient to substantially suppress the climbing deficit and mitochondrial disruption in *Pink1* and *parkin* mutants (Fig. 6A-D), as previously reported (18). Remarkably, coincident knockdown of *fwd* completely prevented the ability of Drp1 to rescue the *Pink1/parkin* mutant phenotypes (Fig. 6A-D). These results further indicates that Drp1 requires the activity of Fwd.

**Figure 6.**
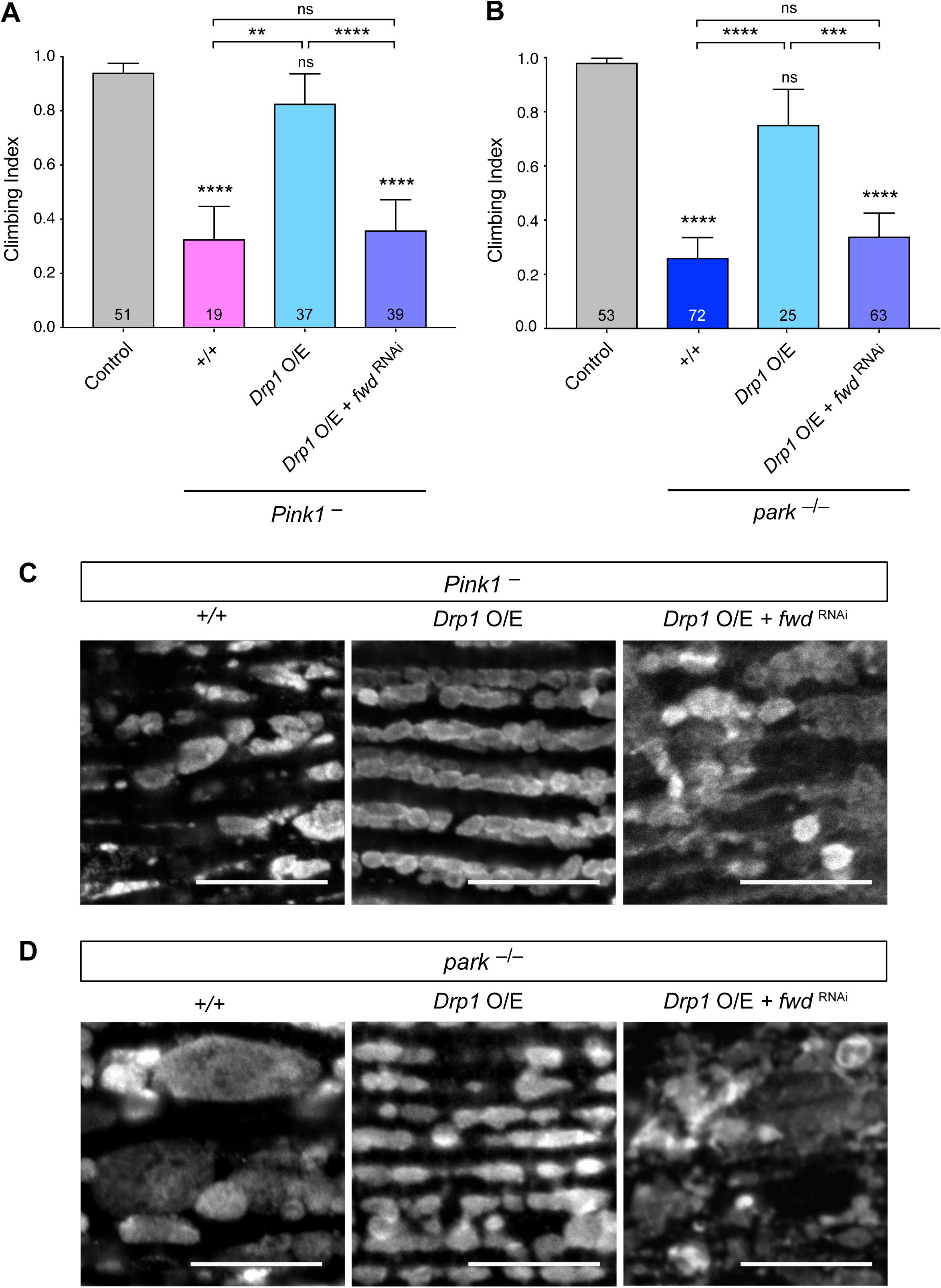
*Drp1* activity requires *fwd* in suppressing *Pink1/parkin* phenotypes. (A, B) Climbing assay of control, *Pink1* or *parkin* mutants with or without *Drp1* overexpression or concomitant induction of *fwd* RNAi. Significance was analysed by Kruskal-Wallis test with Dunn’s post-hoc correction for multiple comparisons. Comparison is against the control unless otherwise indicated; ** *P<*0.01, *** *P<*0.001, **** *P<*0.0001; ns, non-significant. (C, D) Confocal microscopy analysis of mitochondrial integrity, visualised by anti-ATP5A immunostaining, in flight muscles of the indicated genotypes. For all conditions, transgenic expression was mediated via *da-GAL4*. Scale bar = 10 μm. Full genotypes are given in Supplementary Table 1.

## Discussion

We previously identified knockdown of *fwd* to induce mitochondrial hyperfusion in cultured cells, similar to loss of *Pink1* (24). Here we have validated that genetic loss or knockdown of *fwd* also causes excess mitochondrial fusion in neuronal cells *in vivo*, leading to increased mitochondrial length and branching (Fig. 2). As mitochondrial fission/fusion dynamics have been shown to be important for proper mitochondrial homeostasis (2), it is not surprising that this also has an impact on respiration at the organismal level (Fig. 3). Furthermore, it follows that this in turn has an impact on organismal fitness and vitality (Fig. 1). While *fwd* mutants have previously been shown to be male sterile, we describe for the first time new phenotypes associated with loss of *fwd*: profound locomotor deficits and shortened lifespan. Interestingly, there is a stronger requirement for *fwd* in the nervous system compared to the musculature.

The robust locomotor phenotype allowed us to test the genetic relationship between *fwd* and core components of the mitochondrial fission/fusion machinery. Given the excess mitochondrial fusion in *fwd* mutants, suppression of the organismal phenotypes by reduction of fusion factors *Marf* and *Opa1* was expected. However, it was surprising that overexpression of the fission factor *Drp1* was unable to ameliorate organismal phenotypes or even the mitochondrial morphology (Fig. 4). These results suggested that Drp1 requires Fwd to function. Consistent with this, *Drp1* overexpression was no longer able to rescue *Pink1/parkin* mutant phenotypes in the absence of *fwd* (Fig. 6). These genetic experiments strongly hint at a functional link between Drp1 and Fwd but do not illuminate the molecular mechanism underpinning it. Fwd, is the *Drosophila* homologue of phosphatidylinositol 4-kinase IIIβ [PI(4)KB], which mediates the phosphorylation of phosphatidylinositol to generate phosphatidylinositol 4-phosphate [PI(4)P] (31). PI(4)P is one of the most abundant phosphoinositides, which is usually concentrated in the trans-Golgi network (32); thus, an obvious mechanism by which PI(4)P may influence mitochondrial dynamics is not immediately apparent. However, while this manuscript was in preparation, Nagashima and colleagues reported that Golgi-derived PI(4)P-containing vesicles were required for the final stages of mitochondrial fission (33). In that study, the authors found that loss of PI(4)KIIIβ led to a hyperfused and branched mitochondrial network, consistent with what we observed here (Fig. 2). Moreover, they described that while Drp1 was still recruited, it was unable to fully execute the scission event, although the reason is unclear, leading to extended mitochondrial constriction sites. Our genetic evidence that the action of Drp1 requires Fwd is consistent with these findings, and provide an *in vivo* validation of Nagashima and colleagues’ results. Further, it is interesting to note that while the study by Nagashima et al. suggests a universal role for PI(4)P in mitochondrial fission, our *in vivo* analysis revealed that while *fwd* affected mitochondrial morphology in the nervous system, it appeared to have no major impact in the musculature. These tissue-specific requirements were borne out in the strong locomotor deficits caused by neuronal loss of *fwd* but much less so by knockdown in muscles. Clearly, further work is required to better understand the complexities of regulated fission/fusion events in different cell contexts *in vivo*.

A key role of mitochondrial fission/fusion dynamics is in contributing to a quality control mechanism of mitochondrial sorting to eliminate dysfunctional units via mitophagy (3, 4). A substantial body of evidence from cellular models indicates that mammalian PINK1/Parkin act to promote damage-induced mitophagy (5-7), and some *in vivo* evidence from *Drosophila* also supports this (34, 35). However, the precise nature of PINK1/Parkin-mediated mitochondrial turnover *in vivo* is debated with contradictory results emerging (36-40). Nevertheless, interventions to combat the decline in mitochondrial homeostasis remain a key challenge to combatting *PINK1/PRKN* related pathologies. One mechanism that seems to provide substantial benefit in physiological contexts is through augmenting mitochondrial fission, which presumably facilitates the flux of damaged mitochondrial components towards turnover (17-20). Here, we provide further evidence that augmenting a pro-fission pathway is beneficial against *Pink1/parkin* dysfunction. As phosphoinositides can be interconverted by the action of multiple enzymes that may be druggable, these findings suggest another potential route towards a therapeutic intervention.

## Methods

### *Drosophila* stocks and husbandry

Flies were raised and kept under standard conditions in a temperature-controlled incubator with a 12h:12h light:dark cycle at 25 °C and 65% relative humidity, on food consisting of agar, cornmeal, molasses, propionic acid and yeast. The following strains were obtained from the Bloomington *Drosophila* Stock Center (RRID:SCR_006457): *w*^1118^ (RRID:BDSC_6326), *fwd*^neo1^ (RRID:BDSC_10069), *Df(3L)7C* (RRID:BDSC_5837), *Opa1*^s3475^ (RRID:BDSC_12188), *da-GAL4* (RRID:BDSC_55850), *nSyb-GAL4* (RRID:BDSC_51941), *Mef2-GAL4* (RRID:BDSC_ 27390), *CCAP-GAL4* (RRID:BDSC_25685, RRID:BDSC_25686), *UAS-mito-HA-GFP* (RRID:BDSC_8442, RRID:BDSC_8443), *fwd*^RNAi^ (RRID:BDSC_35257), *luciferase*^RNAi^ (RRID:BDSC_31603). Other lines were kindly provided as follows: *fwd*^3^ from J. Brill (27), and the *Pink1*^B9^ and UAS-*Drp1* from J. Chung (10), *Marf*^B^ from H. Bellen (41). The *park*^25^ mutants have been described previously (8). UAS-*GFP- fwd* was generated by PCR amplification of the GFP-fwd sequence from a *hsp83::GFP-fwd* plasmid (28), kindly provided by G. Polevoy and J. Brill, and cloned into pUAST.attB for integration at the attP40 locus (BestGene Inc.). All experiments in adult flies were conducted using males, except Fig. 4A where females were used.

### Locomotor assays

The startle induced negative geotaxis (climbing) assay was performed using a counter-current apparatus. Experiments were performed using 2-3 days old flies. Expect for figure 4A, all the climbing assays used males. Briefly, 20-23 flies were placed into the first chamber, tapped to the bottom, and given 10 s to climb a 10 cm distance. This procedure was repeated five times (five chambers), and the number of flies that has remained into each chamber counted. The weighted performance of several group of flies for each genotype was normalized to the maximum possible score and expressed as *Climbing index* (8).

### Lifespan

For lifespan experiments, flies were grown under identical conditions at low density. Progeny were collected under very light anaesthesia and kept in tubes of approximately 25 males each, and transferred every 2-3 days to fresh media and the number of dead flies recorded. Percent survival was calculated at the end of the experiment after correcting for any accidental loss.

### Immunohistochemistry and sample preparation

For immunostaining, adult flight muscles were dissected in PBS and fixed in 4% formaldehyde for 30 min at RT, permeabilized in 0.3% Triton X-100 for 30 min, and blocked with 0.3% Triton X-100 plus 4% Horse Serum (HS) in PBS for 1 h at RT. Tissues were incubated with ATP5A antibody (Abcam Cat# ab14748, RRID:AB_301447; 1:500), diluted in 0.3% Triton X-100 plus 4% HS in PBS overnight at 4°C, then rinsed 3 times 10 min with 0.3% Triton X-100 in PBS, and incubated with the appropriate fluorescent secondary antibodies overnight at 4°C. The tissues were washed 2 times in PBS and mounted on slides. Adult brains were dissected in PBS and fixed on ice in 4% formaldehyde for 30 min, permeabilized in 0.3% Triton X-100 for 30 min, and blocked with 0.3% Triton X-100 plus 4% HS in PBS for 4 h at RT. Incubation with Tyrosine Hydroxylase Antibody (TH) antibody (Inmunostar Cat#22941, 1:200) diluted in 0.3% Triton X-100 plus 4% HS was done for 72h at 4°C. Secondary antibody was incubated for 3 h at RT. Then washes were done 3 times for 20 min with 0.3% Triton X-100 in PBS and mounted in carved slides. Larvae brains were dissected on PBS and mounted sideways on slides coated with poly-lysine at 0.9 mg/ml. They were fixed in 4% formaldehyde for 20 min at RT, then washed in PBS. All the sample preparations were mounted using Prolong Diamond Antifade mounting medium (Thermo Fischer Scientific Cat# P36961).

### Microscopy

Fluorescence imaging was conducted using a Zeiss LSM 880 confocal microscope (Carl Zeiss MicroImaging) equipped with Nikon Plan-Apochromat 100x/1.4 NA oil immersion objectives. Images were taken at a resolution of 2048×2048 pixels and they were prepared using Fiji software (RRID:SCR_002285).

### Analysis of mitochondrial morphology

Motoneuron cell bodies from larvae ventral nerve cord expressing *CCAP*-*GAL4* were used to analyse mitochondrial branches marked by mitoGFP. All images were processed using Fiji software (RRID:SCR_002285). Z-stacks of individual neurons were cropped to a size of 232 × 232 pixels. The mitoGFP signal was enhanced and smoothed using two filters: unsharp mask (radius =10.0 pixels, Mask strength 0.9) and median filtering (radius =3). Then binary masks were created using “Otsu method” in auto and Dark background, and ‘skeletonized’ from the Process and Binary menu served to generate the branches. These skeletonized images were analysed using Analyse Skeleton (2D/3D). Finally, Median branch length per cell was calculated using Branch Length column from “Branch information” window, and the proportion of individual vs interconnected branches per cell was calculated by taking the “Number of branches” column from the “Results” window.

### Transmission electron-microscopy

Thoraces were prepared from 5-day-old adult flies and treated as previously described (8). Ultra- thin sections were examined using a FEI Tecnai G2 Spirit 120KV transmission electron-microscope.

### Respirometry analysis

Respiration was monitored at 30 °C using an Oxygraph-2k high-resolution respirometer (OROBOROS Instruments) using a chamber volume set to 2 mL. Calibration with air-saturated medium was performed daily. Data acquisition and analysis were carried out using Datlab software (OROBOROS Instruments). Five flies per genotype (equal weight) were homogenised in respiration buffer (120 mM sucrose, 50 mM KCl, 20 mM Tris-HCl, 4 mM KH_2_PO_4_, 2 mM MgCl_2_, and 1 mM EGTA, 1 g/l fatty acid-free BSA, pH 7.2). For coupled (state 3) assays, complex I-linked respiration was measured at saturating concentrations of malate (2 mM), glutamate (10 mM) and adenosine diphosphate (ADP, 2.5 mM). Complex II-linked respiration was assayed in respiration buffer supplemented with 0.15 µM rotenone, 10 mM succinate and 2.5 mM ADP. Data from 7-8 independent experiments were averaged.

### ATP levels

The ATP assay was performed as described previously (24). Briefly, five male flies for each genotype were homogenized in 100 μL 6 M guanidine-Tris/EDTA extraction buffer and subjected to rapid freezing in liquid nitrogen. Homogenates were diluted 1/100 with the extraction buffer and mixed with the luminescent solution (CellTiter-Glo Luminescent Cell Viability Assay, Promega). Luminescence was measured with a SpectraMax Gemini XPS luminometer (Molecular Devices). The average luminescent signal from technical triplicates was expressed relative to protein levels, quantified using the Pierce BCA Protein Assay kit (ThermoFisher Scientific). Data from 3 independent experiments were averaged and the luminescence expressed as a percentage of the control.

### Statistical analysis

Data from the various experimental assays were analysed as follows: For behavioural analyses, Kruskal-Wallis non-parametric test with Dunn’s post-hoc correction for multiple comparisons was used. Lifespan was analysed by Log-rank (Mantel-Cox) test. Categorical analyses (i.e. thoracic indentations) were analysed by Chi-square test. Mitochondrial branch length by Mann-Whitney non-parametric test, and connectivity by Kruskal-Wallis with Dunn’s post-hoc correction. ATP levels were analysed by unpaired *t*-test, and respiration by paired *t*-test. Analyses were performed using GraphPad Prism 8 software (RRID:SCR_002798) and RStudio software (RRID:SCR_000432).

## Data availability

All data that support the findings of this study are available on reasonable request to the corresponding author. The contributing authors declare that all relevant data are included in the paper.

## Acknowledgements

This work is supported by MRC core funding (MC_UU_00015/4, MC-A070-5PSB0 and MC_UU_00015/6) and ERC Starting grant (DYNAMITO; 309742). Stocks were obtained from the Bloomington *Drosophila* Stock Center which is supported by grant NIH P40OD018537. We thank J. Brill for sharing *fwd* stocks, Claire Pilgrim for technical assistance, and J. Prudent for sharing results prior to publication. We thank members of the Whitworth lab for fruitful discussions and feedback on the manuscript.

## Author Contributions

A.T-F. and E.L.W. designed and performed experiments, and analysed data. A.J.W. conceived the study, designed experiments, analysed the data and supervised the work. A.J.W. wrote the manuscript with input from all authors.

---

**Supplementary Table 1.**
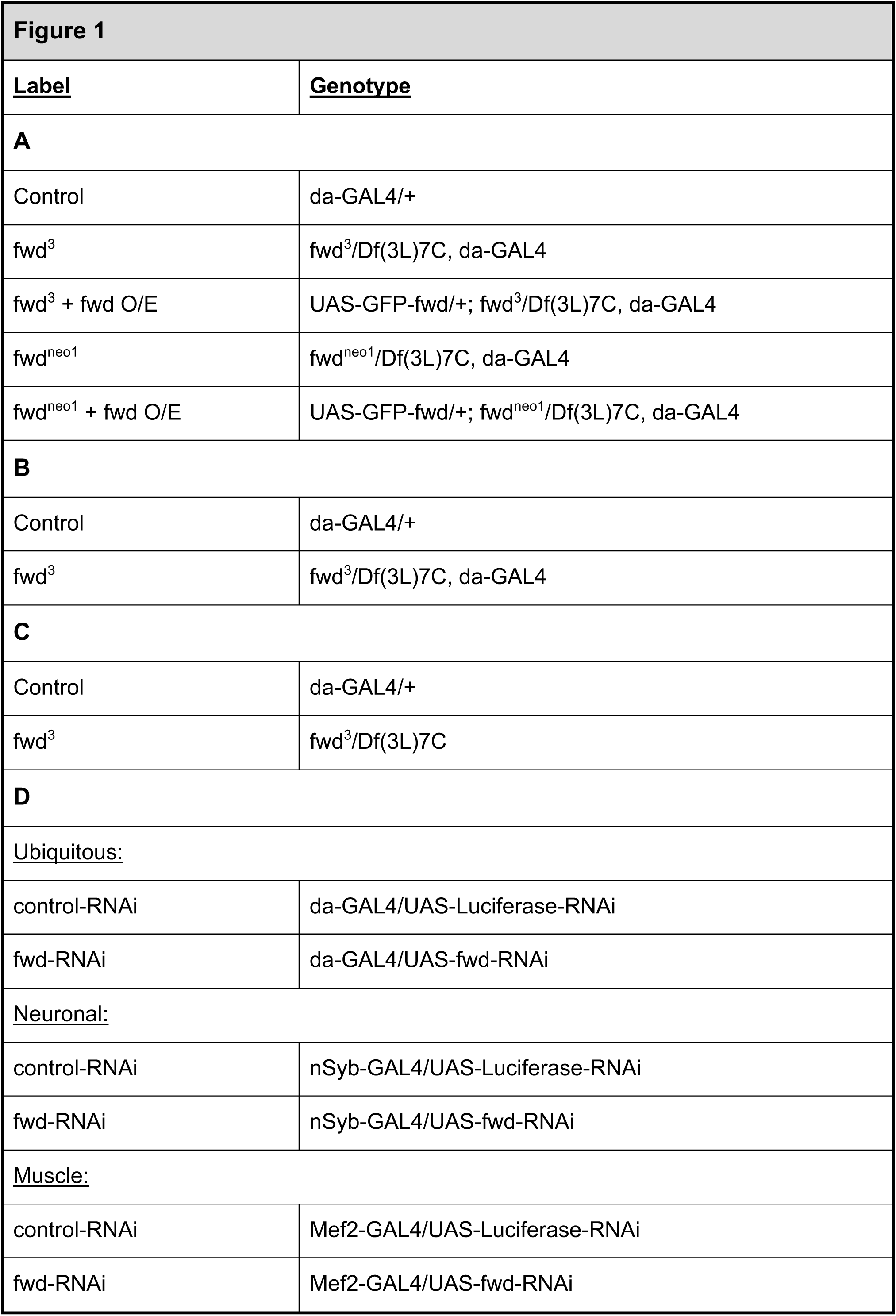
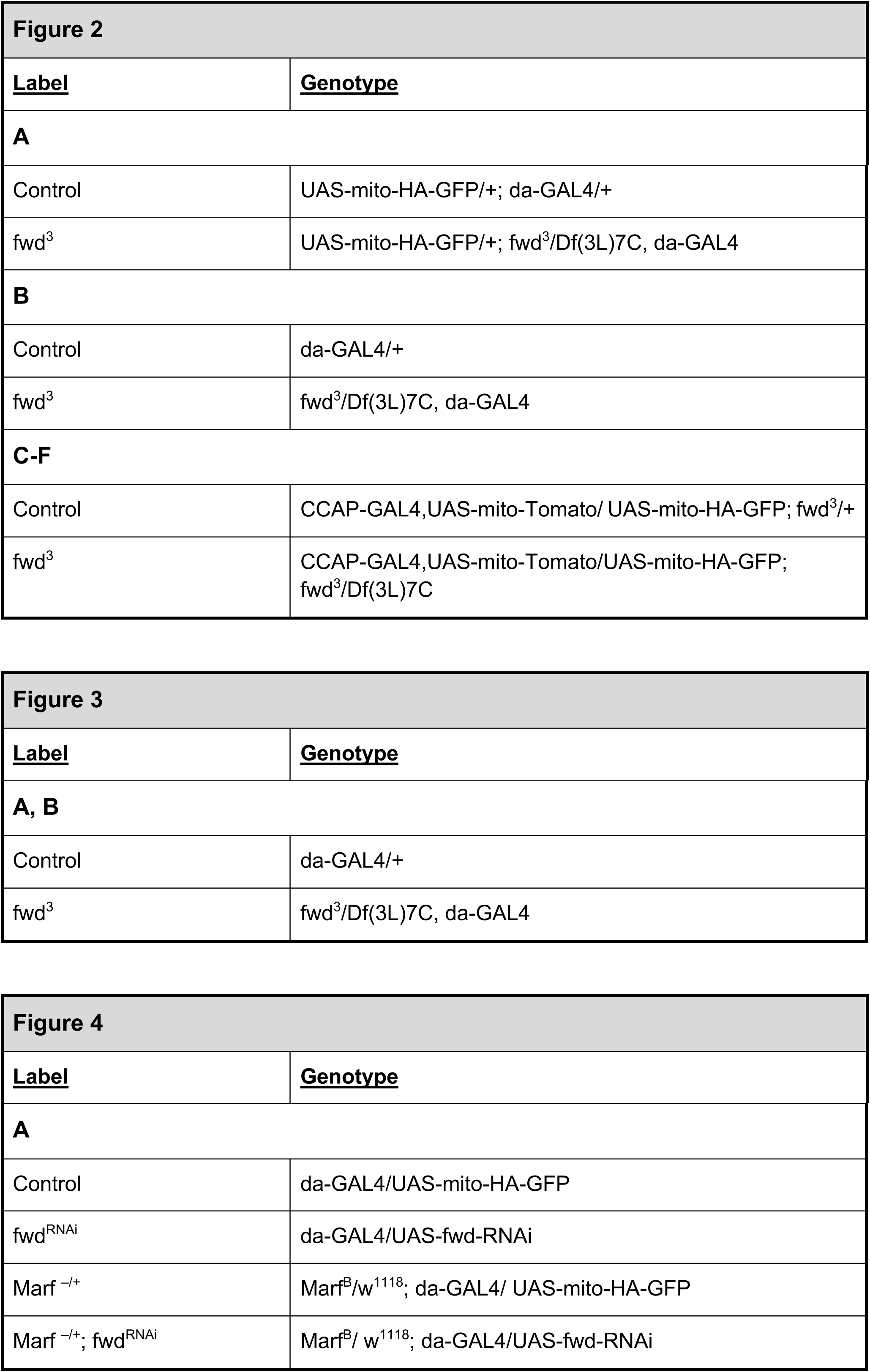
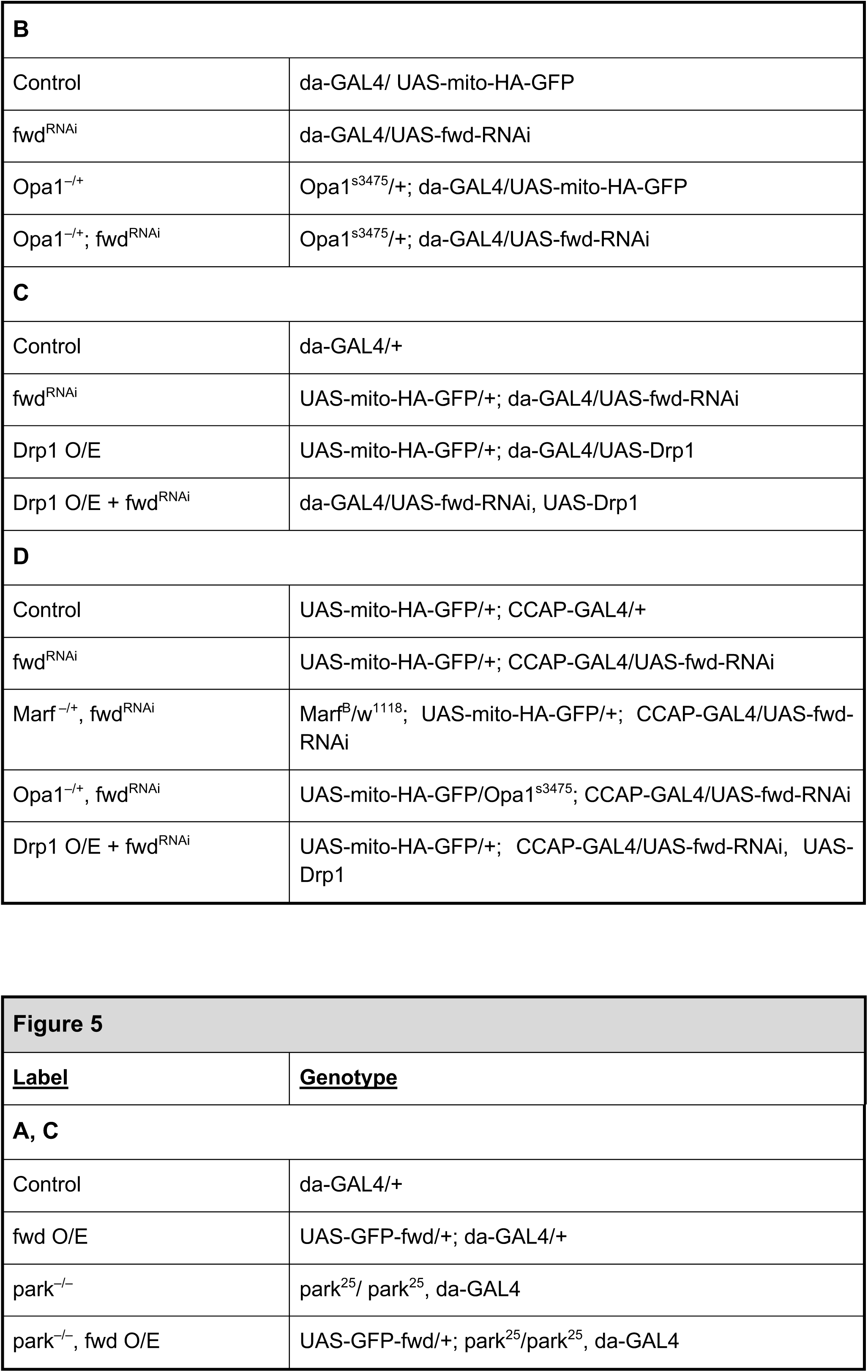
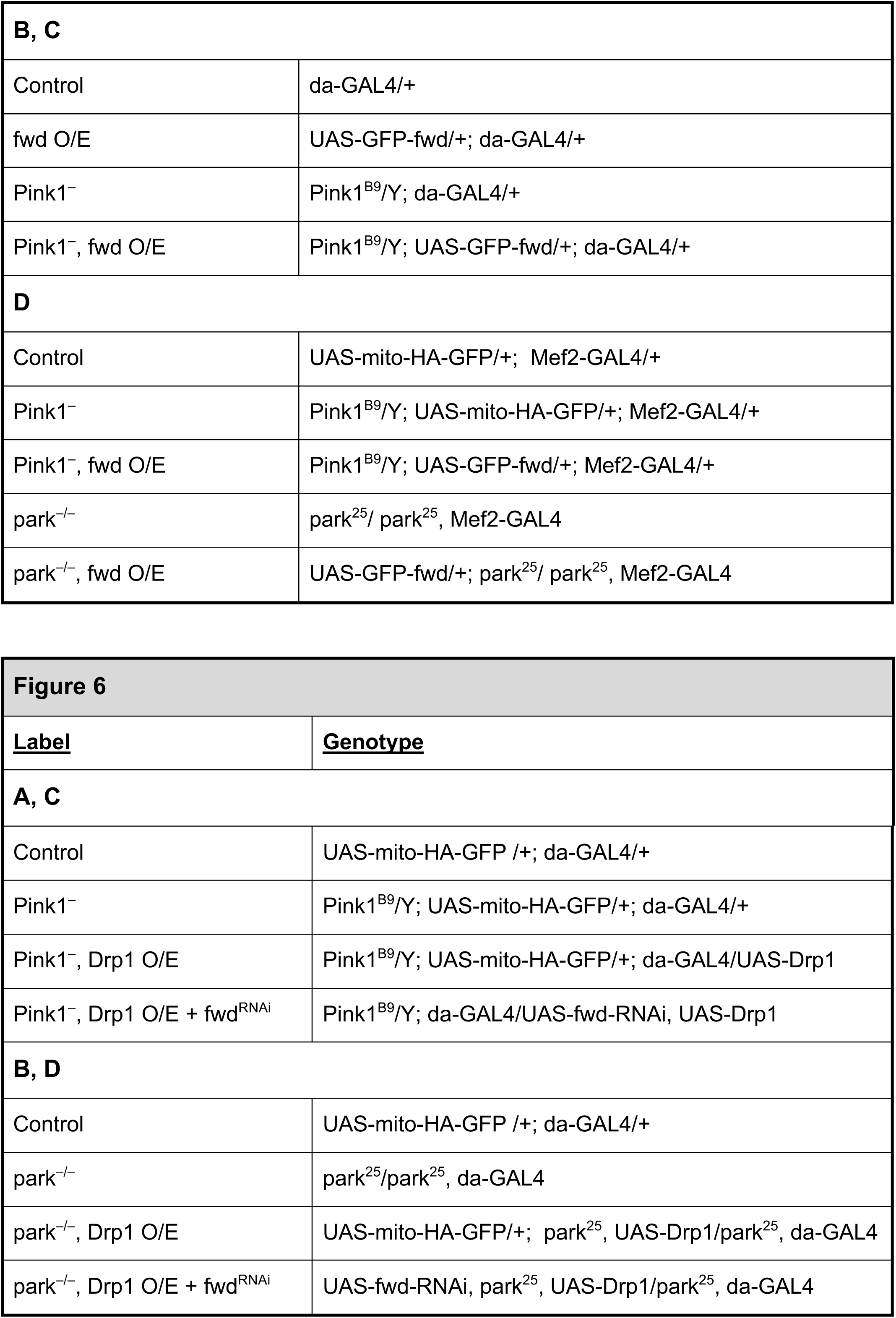
Details of full genotypes used in this study. More details of each line can be found in Methods.

